# Neuronal gamma-band synchronization regulated by instantaneous modulations of the oscillation frequency

**DOI:** 10.1101/070672

**Authors:** E. Lowet, M. J. Roberts, A. Peter, B. Gips, P. De Weerd

## Abstract

Neuronal gamma-band synchronization shapes information flow during sensory and cognitive processing. A common view is that a stable and shared frequency over time is required for robust and functional synchronization. To the contrary, we found that non-stationary instantaneous frequency modulations were essential for synchronization. First, we recorded gamma rhythms in monkey visual area V1, and found that they synchronized by continuously modulating their frequency difference in a phase-dependent manner. The frequency modulation properties regulated both the phase-locking and the preferred phase-relation between gamma rhythms. Second, our experimental observations were in agreement with a biophysical model of gamma rhythms and were accurately predicted by the theory of weakly coupled oscillators revealing the underlying theoretical principles that govern gamma synchronization. Thus, synchronization through instantaneous frequency modulations represents a fundamental principle of gamma-band neural coordination that is likely generalizable to other brain rhythms.

## INTRODUCTION

Synchronization, the ability of oscillators to mutually adapt their rhythms (Pikovsky et al., 2002; Winfree, 1967), is a ubiquitous natural phenomenon. Neural synchronization in the gamma-range (25-80Hz) has been reported both in subcortical structures (Akam et al., 2012; Steriade et al., 1993; Zhou et al., 2016) and in cortical areas (Fries, 2015; Gray and Singer, 1989; Gregoriou et al., 2009). Gamma rhythms emerge in activated neural circuits, in which fast-spiking inhibitory neurons play a central role (Cardin et al., 2009; Tiesinga and Sejnowski, 2009; Traub et al., 1996). A prime example is the emergence of gamma rhythms in the early visual cortex during visual stimulus processing (Brunet et al., 2013; Gail et al., 2000; Gray and Singer, 1989; Hermes et al., 2014; Ray and Maunsell, 2010; Roberts et al., 2013). Gamma synchronization has been found to relate to the formation of neural assemblies within (Gail et al., 2000; Gray and Singer, 1989; Havenith et al., 2011; Vinck et al., 2010) and across brain areas (Bosman et al., 2012; Gregoriou et al., 2009; Jia et al., 2013a; Roberts et al., 2013; Sirota et al., 2008; Zhou et al., 2016). Precise temporal coordination of presynaptic spikes increases their effectiveness on postsynaptic targets (Fries et al., 2001; Tiesinga et al., 2005) and can thereby modulate the effectiveness of neural communication (Börgers et al., 2005; Cannon et al., 2014; Womelsdorf et al., 2007), as shown between V1 and V4 during visual attention (Bosman et al., 2012; Grothe et al., 2012). Temporal coordination in terms of spike timing (phase code) might be an efficient and robust mechanism for information coding (Havenith et al., 2011; Jensen et al., 2014; Maris et al., 2016; Tiesinga et al., 2008; Vinck et al., 2010). Further, gamma rhythmic inhibition might increase coding efficiency through sparsening (Chalk et al., 2015; Jadi and Sejnowski, 2014; Vinck and Bosman, 2016) and normalization (Gieselmann and Thiele, 2008; Ray et al., 2013) of neural activity. These network consequences of gamma have led to influential hypotheses about the function of gamma for sensation and cognition (Buehlmann and Deco, 2010; Buzsáki and Wang, 2012; Eckhorn et al., 2001; Fries, 2015; Gray and Singer, 1989; Maris et al., 2016; Miller and Buschman, 2013), including a role in perceptual grouping (Eckhorn et al., 2001; Engel et al., 1999; Gray and Singer, 1989) and in visual attention (Bosman et al., 2012; Fries, 2015; Gregoriou et al., 2009; Miller and Buschman, 2013).

Surprisingly, in spite of important scientific advances, it is not well understood how gamma rhythms synchronize and what the underlying principles of synchronization are. For example, recent experimental observations of large variability of the precise oscillation frequency have raised doubts on the robustness and functionality of gamma synchronization in the brain. It has been observed that the precise frequency fluctuates strongly over time (Atallah and Scanziani, 2009; Burns et al., 2011, 2010) and that different cortical locations can express different preferred frequencies (Bosman et al., 2012; Ray and Maunsell, 2010). That these observations have led to doubts on the functionality of gamma synchronization indicates that research into gamma synchronization often starts from the premise that continuously matched frequencies are a requirement for the occurrence of stable phase-relations. The observation of frequency variations and frequency differences would then suggest that meaningful synchronization cannot be maintained. These ideas reveal a stationary view of synchronization, which assumes that the underlying oscillatory dynamics are stable at a fixed phase-relation and shared frequency. This is also reflected in the widespread use of stationary methods to assess gamma synchronization, of which spectral coherence is a prime example (Carter et al., 1973). From a dynamic systems perspective however, synchronization is primarily a non-stationary process (Izhikevich and Kuramoto, 2006; Izhikevich, 2007; Kopell and Ermentrout, 2002; Pikovsky et al., 2002; Winfree, 1967), because oscillators adjust their rhythms through phase shifts (i.e., changes in the instantaneous frequency).

Here, by using a combination of theoretical and experimental techniques, we studied the dynamical principles of gamma synchronization in monkey visual area V1. We simultaneously recorded gamma-rhythmic neural activity at different V1 cortical locations and studied their synchronization properties while using local stimulus contrast (Ray and Maunsell, 2010) to modulate the frequency difference (detuning). Strikingly, we observed that frequency-variable gamma rhythms still synchronized, even when the mean frequencies did not match. This was achieved by continuously varying their instantaneous frequency difference in a manner depending on the phase difference. The function relating phase difference to frequency difference had a sinusoidal-like shape. The interplay between the detuning, representing a desynchronization force, and the amount of instantaneous frequency modulations, representing a synchronization force, regulated the phase-locking strength and the preferred phase-relation between V1 locations. Further, detuning was dependent on visual grating contrast difference, whereas frequency modulation strength was dependent on the cortical distance.

To assess the biophysical underpinning of our V1 observations, we simulated two interacting pyramidal-interneuron gamma (PING) networks (Bartos et al., 2007; Börgers et al., 2005; Tiesinga and Sejnowski, 2009). In line with our observation in V1, we found gamma synchronization to be associated with rapid frequency modulations. The modulation strength was modulated by synaptic connectivity, whereas detuning was dependent on the excitatory input drive. To achieve a principled understanding of our observations, we applied the theoretical framework of weakly coupled oscillators (Ermentrout and Kleinfeld, 2001; Hoppensteadt and Izhikevich, 1998; Kopell and Ermentrout, 2002; Kuramoto, 1991; Pikovsky et al., 2002). We found that a single differential equation accounted well for the non-stationary frequency modulations and further allowed for precise predictions of how the phase-locking and the phase-relation between gamma rhythms changed across conditions.

## RESULTS

### Local frequency differences regulate the dynamic synchronization process between V1 gamma rhythms

We first asked how synchronization within V1 was influenced by frequency differences, and by the distance between recording sites. To this aim, we recorded from 2 to 3 laminar probes simultaneously in cortical area V1 of two macaques (M1 and M2) (Fig.1A). We used distances in the order of magnitude of V1 horizontal connectivity (Stettler et al., 2002), hence probes were separated by 1 to 6mm. Using laminar probes enabled us to reduce the influence of volume conduction by calculating current-source density (CSD) as a network signal. Using CSD, we estimated the instantaneous frequency, phase and phase difference of gamma signals. The monkeys fixated centrally while a whole-field static grating with spatially variable contrast was shown. Gamma power was induced in layers 2-4 and in the deepest layer (Fig.1B, Fig.S1). V1 locations showed increased gamma frequency with increased local contrast (linear regression, single contact level, M1: R^2^=0.38, M2: R^2^= 0.27, both p<10^−10^, Fig.1C, Fig.S2) allowing us to parametrically vary the frequency difference between probes by varying the contrast difference. We will first show the key results through three illustrative examples. In the first example, we chose two cortical locations separated by a relatively large distance of ~5mm, presented with a visual contrast difference of 17% (Fig.1D). Their frequency difference was 5Hz as shown by their non-overlapping power spectra (Fig.1E). This would imply that the phase difference would not be constant, but would advance at a phase precession rate of 2π every 200ms, which could be expected to preclude synchronization. However, the frequency difference was not constant. Instead, the instantaneous frequency difference was modulated as a function of phase difference (Fig.1F, Fig.S3) with a modulation amplitude of 1Hz. At the smallest frequency difference (4Hz, yellow point) the phase precession was slowest, at 2π every 250ms, meaning that the oscillators stayed relatively longer around that phase difference. As a result, the probability distribution of phase differences over time (Fig.1G) was non-uniform giving a phase-locking value (Lachaux et al., 1999) (PLV) of 0.11. The peak of the distribution, the ‘preferred phase’, was at 1.3rad, in line with the minimum of the instantaneous frequency modulation function. In the second example, we chose a pair with a similar frequency difference of 4.8Hz but a closer distance (~2.5mm, Fig.1H). The instantaneous frequency modulation amplitude was larger with a modulation amplitude of 1.8Hz (Fig.1J) and a modulation minimum around 3Hz at the preferred phase. Because phase precession at the preferred phase was slower, the phase difference distribution was narrower than in the previous example, indicating higher synchrony (PLV=0.32, Fig.1K) with a peak centered at a different phase (0.78rad). In the third example the cortical distance remained the same but the frequency difference was reduced (2.8Hz) by eliminating the contrast difference (Fig.1M, the remaining frequency difference might be due to eccentricity, see Fig.S2). The frequency modulation amplitude did not change however, with a lower mean difference, the modulation minimum was close to zero (1Hz, Fig.1N), thus the associated phase difference (0.48rad) could be maintained for even longer periods and the phase difference probability distribution was even narrower (PLV=0.51, Fig.1O). The three examples were representative for the 805 recorded contact pairs in monkey M1 and 882 contact pairs in monkey M2 where each pair was recorded at 9 levels of contrast difference.

**Fig.1.**
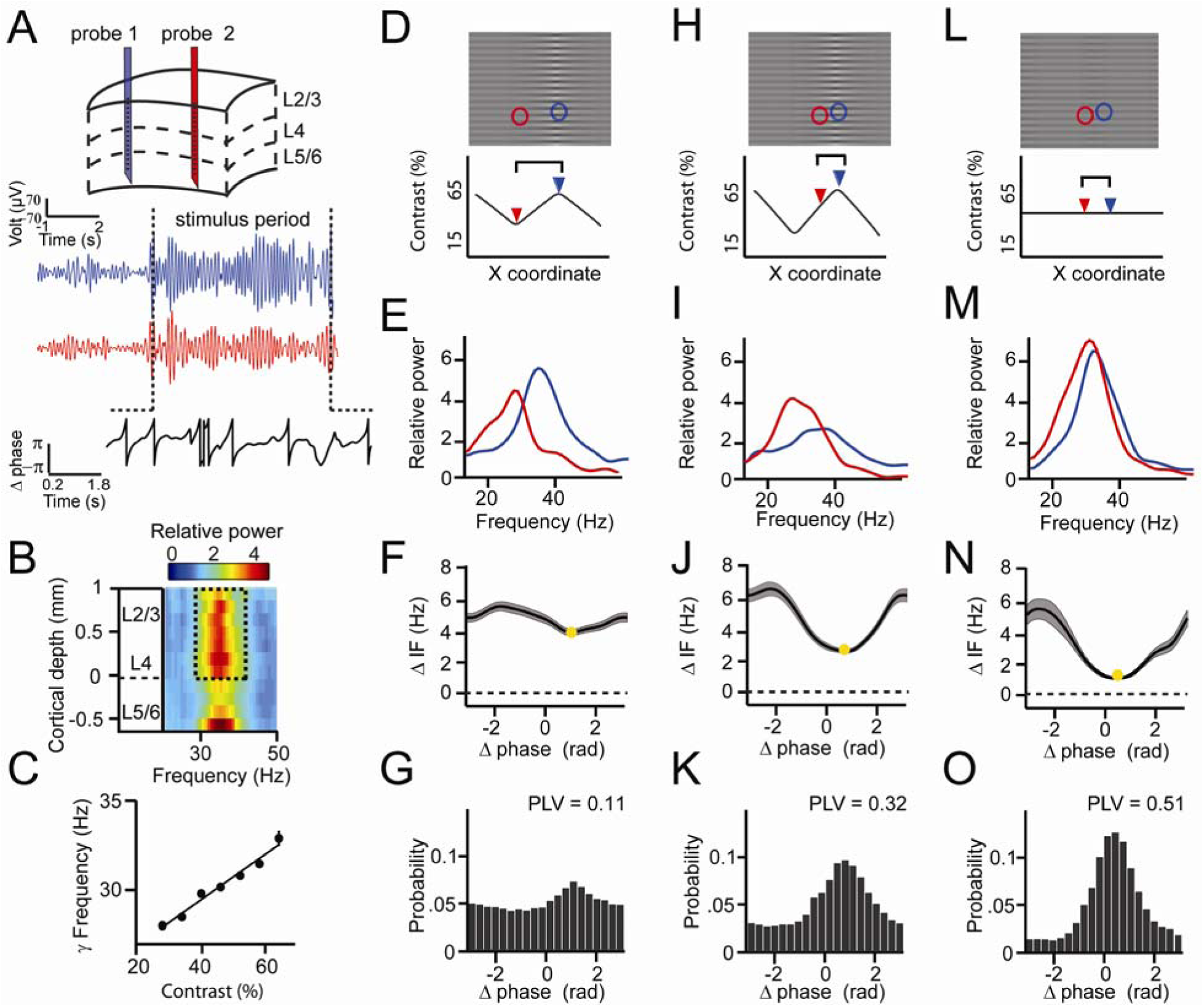
*Experimental paradigm and intermittent synchronization. **(A)** Recordings preparation and example CSD (blue and red) traces from which phase difference (black) trace was extracted. The gradient of the black trace indicates the rate of phase precession. **(B)** Spectral power relative to baseline as a function of V1 cortical depth (36.5% contrast, population average, M1) dashed box indicates gamma in the layers taken for main analysis **(C)** Local contrast modulated gamma frequency (population average, M1). **(D-G)** Example 1 showing synchronization despite frequency difference. **(D**) Section of the stimulus grating. Two receptive fields (RF) from different probes are superimposed (blue and red circles). Below, black line gives contrast over space, arrowheads mark RF positions. **(E)** Power spectra of the two probes showing different peak frequencies. **(F)** Instantaneous frequency difference (ΔIF), equivalent to the phase precession rate, as a function of phase difference. Yellow dot indicates the modulation minimum, equivalent to the preferred phase difference, shading is ±SE (**G)** The phase difference probability distribution and phase-locking value (yellow dot, PLV). **(H-K)** Example 2; probes were closer, gamma peak frequency difference was similar. Conventions as in D-G. **(L-O)** Example 3; same distance, reduced frequency difference. Compare F, J, N; the RF distance determined IF modulation amplitude, whereas contrast difference determined mean gamma frequency difference*

### Experimental observations reproduced by two weakly coupled pyramidal-interneuron gamma (PING) networks

To gain a first understanding of our experimental observations, we tested whether the findings were reproducible by a well-established biophysical model of cortical gamma rhythms (see Supplementary Information for more details). We simulated two coupled pyramidal-interneuron gamma (PING) networks (Fig.2A), which have been shown to capture many properties of cortical gamma rhythms (Börgers et al., 2005; Jadi and Sejnowski, 2014; Lowet et al., 2015; Tiesinga and Sejnowski, 2009, 2010). The network consisted of excitatory regular-spiking spiking neurons, representing pyramidal neurons, and fast-spiking inhibitory interneurons. We used the Izhikevich neural model (Izhikevich, 2003). Neurons were connected through excitatory AMPA and inhibitory GABA-A synapses. To mimic V1 horizontal connections (Stettler et al., 2002), the two PING networks were weakly coupled through excitatory cross-network connections that targeted the excitatory and inhibitory neurons of the receiving network. Each network received an independent source of excitatory drive, mimicking the effect of local visual contrast (Sclar et al., 1990). Neurons also received additional noise, such that the oscillation frequency was instable over time as observed for V1 gamma. For each network we estimated a population signal from which we extracted the instantaneous phase (Fig.2B). In line with our experimental observations and previous studies (Jia et al., 2013b; Lowet et al., 2015; Roberts et al., 2013; Tiesinga and Sejnowski, 2009), the input drive set the frequency of the gamma rhythm (R^2^=0.98, Fig.2C). To reproduce the experimental V1 findings shown in Fig.1 (Fig.2D-O) we modulated the cross-network connection strength, mimicking cortical distance, and the difference of input drive between networks, mimicking the local contrast difference (Fig.2D, H, L). These manipulations led to effects on the spectra (Fig.2E,I,M), on the relationship of instantaneous frequency difference to phase difference (Fig.2F,J,N), and on phase-relation distributions (PLV and preferred phase difference) that were similar to those observed in the empirical V1 data. In particular, the modulation of the frequency difference between the gamma rhythms as a function of phase difference had an approximatively sinusoidal shape in the model data, as in the empirical V1 data. Stronger synchronization of gamma rhythms was associated with larger non-stationary modulations of the frequency difference. The strength of the modulation was changed by the synaptic connectivity between networks, whereas the input drive difference changed the frequency difference. As in V1, the phase difference probability distribution was determined by the frequency difference modulations: The mean frequency difference and the amplitude of the frequency modulation defined both the preferred phase-relation and the narrowness of the distribution (PLV). Taken together, this shows that the observations of V1 gamma can be accurately modelled by mutually interacting PING networks, in which synchronization is shaped by the phase-dependent instantaneous frequency modulations.

**Fig.2.**
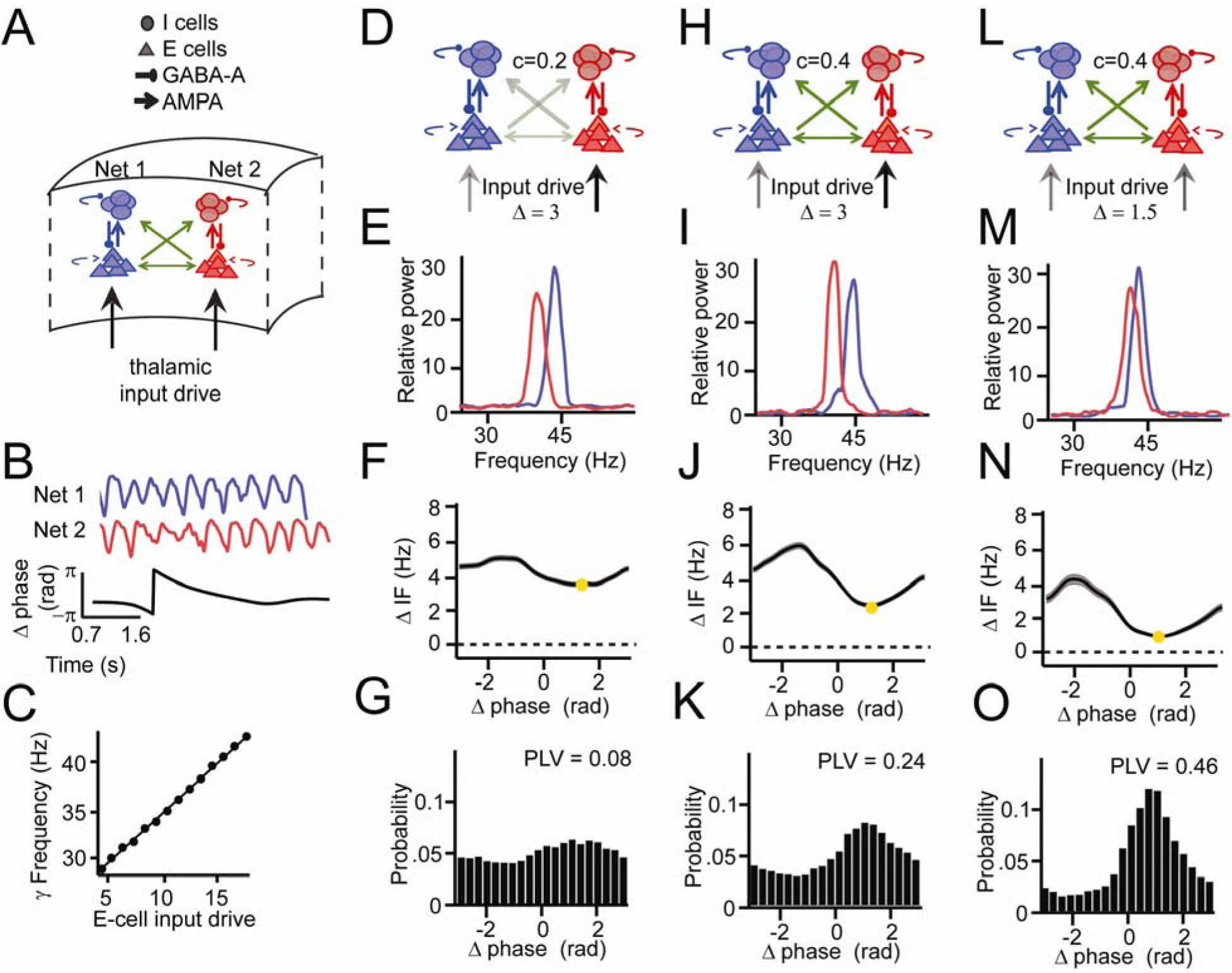
*PING network simulations and intermittent synchronization. **(A)** Two coupled pyramidal-interneuron gamma (PING) networks (Net 1 and Net 2). **(B)** Simulation output example network signals (red and blue) and phase difference θ (black) **(C)** The frequency of gamma in a single network depends on input strength. **(D-G)** Example 1 showing synchronization despite frequency difference. **(D)** Net 1 and Net 2 were relatively weakly coupled (c=0.2, where c defines max synaptic connection strength of a uniform distribution [0,max]) and received a relatively large input difference. **(E)** Power spectra of the two networks showed different peak frequencies. **(F)** Instantaneous frequency difference (ΔIF), equivalent to phase precession rate, as a function of phase difference. Yellow dot indicates the modulation minimum equivalent to the preferred phase difference, shading is ±SE (**G)** The phase difference probability distribution and phase-locking value (PLV). **(H-K)** Example 2; networks were more strongly connected (c=0.4), gamma peak frequency difference was similar. Conventions as in D-G. **(L-O)** Example 3; same connection strength, yet reduced frequency difference. Compare F, J, N; the connection strength determined IF modulation amplitude, whereas input difference determined mean gamma frequency difference.*

### The theory of weakly coupled oscillators (TWCO): A framework for cortical gamma synchronization

We now show how the observed synchronization behavior can be accounted for within the mathematical framework of the theory of weakly coupled oscillators (Ermentrout and Kleinfeld, 2001; Hoppensteadt and Izhikevich, 1998; Kopell and Ermentrout, 2002; Kuramoto, 1991; Pikovsky et al., 2002; Winfree, 1967). Many oscillatory phenomena in the natural world represent dynamic systems with a limit-cycle attractor (Winfree, 2001). Although the underlying system might be complex (e.g. a neuron or neural population), the dynamics of the system can be reduced to a phase-variable if the interaction among oscillators is weak. If interaction strength is weak, amplitude changes are relatively small and play a minor role in the oscillatory dynamics. In this way, V1 neural populations can be approximated as oscillators, ‘weakly coupled’ by horizontal connections. The manner in which mutually coupled oscillators adjust their phases, by phase-delay and phase-advancement, is described by the phase response curve, the PRC (Brown et al., 2004; Canavier, 2015; Izhikevich, 2007; Kopell and Ermentrout, 2002; Schwemmer and Lewis, 2012). The PRC is important, because if the PRC of a system can be described, the synchronization behavior can be understood at a more general level and hence predicted across various conditions.

According to the theory, the synchronization of two coupled oscillators can be predicted from the forces they exert on each other as a function of their instantaneous phase difference. The amount of force is here defined as interaction strength and the interaction function as the PRC. Each oscillator has an intrinsic (natural) frequency and additionally an own source of phase noise, making the oscillators stochastic. The phase precession of two oscillators is given by (Fig.3A):

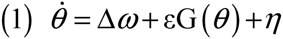

where 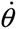 is the time derivative of the phase difference *θ* (the rate of phase precession), Δ*ω* the detuning (the intrinsic frequency difference), ε the interaction strength, G(*θ*) is defined as the mutual PRC, and □ the combined phase noise, where 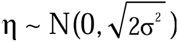. Phase noise is defined here as variation, unrelated to interaction, that occurs for neural oscillators due to inherent instabilities of the generation mechanism (Atallah and Scanziani, 2009; Burns et al., 2010). This type of variation is distinct from measurement noise that is unrelated to the dynamics of the system. We express *ω*, ε and □ in units of Hz (1Hz=2π*rad/s). The time derivative 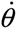 is also expressed in Hz (instantaneous frequency, IF). The equation was solved analytically (see Supplementary Information) to study changes in the phase-difference probability distribution, here characterized by the PLV and the mean (preferred) phase difference, as a function of detuning Δ*ω* and interaction strength ε. The model’s behavior as a function of detuning Δ*ω* and interaction strength ε can be understood more easily by considering the noise-free case first. In the noise-free case (σ=0) one can solve the equation for zero-points (equilibrium points), meaning that the phase precession is zero, (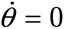, i.e. zero frequency difference). To reach equilibrium, the detuning Δω and the interaction term εG(θ) need to be counterbalanced. When detuning is smaller than the interaction strength (Δ*ω*<=ε), then there is a particular phase difference θ at which an equilibrium can be reached. At equilibrium, there is no phase precession (Fig.3B) and thus a PLV of 1 (full synchronization). When interaction strength is zero (ε=0), the asynchronous oscillators display continuous linear phase precession and have zero PLV (Fig.3C), with the exception of zero detuning. When detuning is larger than a nonzero interaction strength (Δ*ω*>ε, ε>0), oscillators exhibit a nonlinear phase precession over time, characteristic for the intermittent synchronization regime (Izhikevich, 2007; Pikovsky et al., 2002, Fig.3D). The phase precession rate (instantaneous frequency difference) is determined by the detuning Δω, the modulation shape G(θ), and the modulation amplitude ε. Around the preferred phase-relation, the instantaneous frequency difference is reduced (‘slow’ precession in Fig.3D), whereas away from the preferred phase-relation, the instantaneous frequency is larger (‘fast’ precession in Fig.3D). In this regime, PLV between 0 and 1 can be obtained. Including phase noise (σ>0) has important effects on the synchronization behavior (Izhikevich, 2007; Pikovsky et al., 2002). The noise flattens the phase-relation distribution and can induce full cycles of phase precession (phase slips) that also lead to instantaneous frequency modulations. Hence, for noisy oscillators, the intermittent synchronization regime is the default regime for a large parameter range.

**Fig.3.**
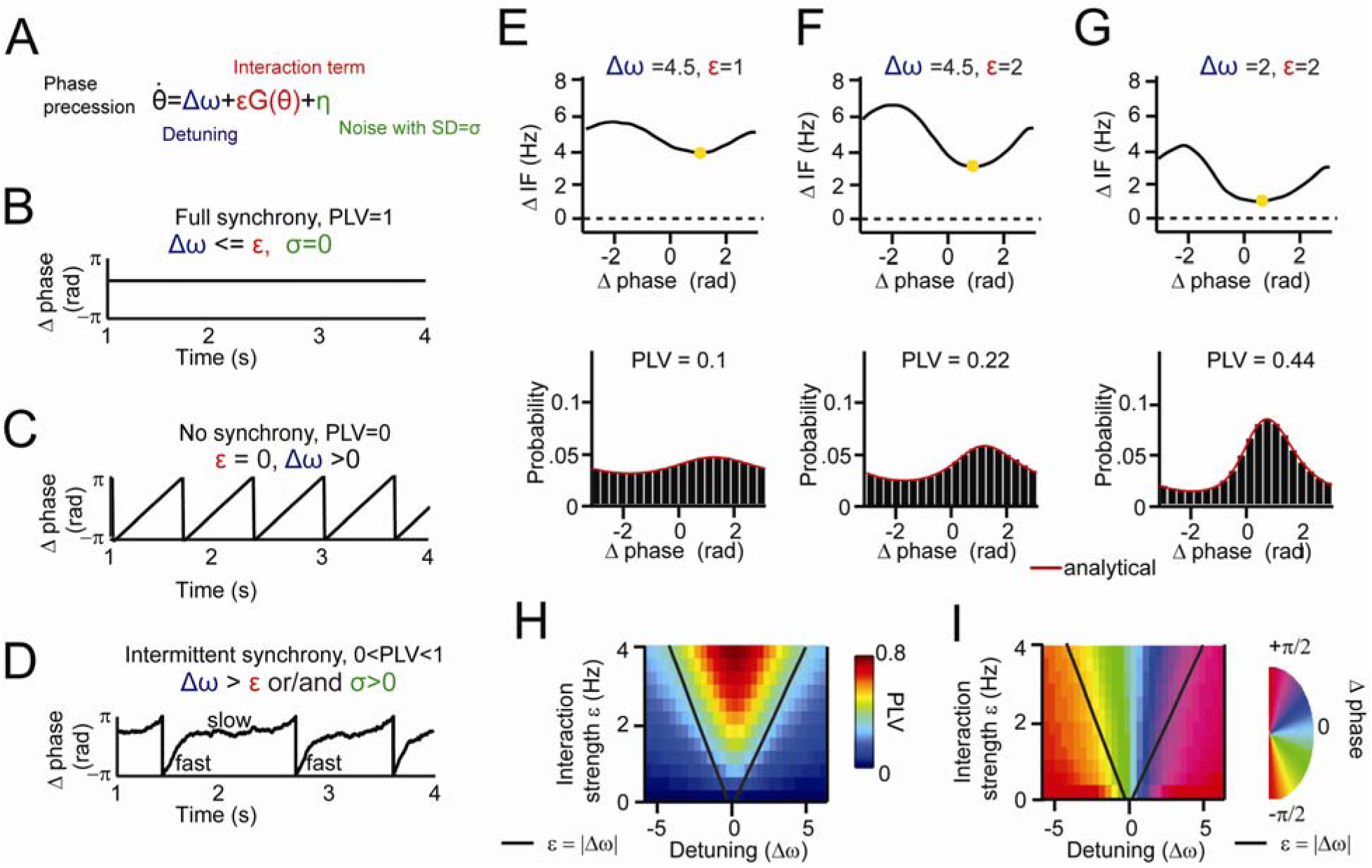
*Theory of weakly coupled oscillators (TWCO). **(A)** The single differential equation used for analysis, with colors representing different key parameters. **(B-D)** Rate of phase precession plotted in different synchronization regimes, with (**B**) full synchrony, (**C**) no synchrony and (**D**) intermittent synchrony. For each plot, the corresponding range of the parameters and the PLV are indicated. **(E-G)** Equivalent behavior as in the examples as Fig.1 and 2. Top is the modulation of the instantaneous frequency difference as a function of phase difference. Bottom is the phase difference probability distribution. Black bars are numerical simulation results, whereas the red line indicates the analytical solution. (**E**) Large detuning and low interaction strength. (**F**) Large detuning and strong interaction strength. (**G**) Small detuning and strong interaction strength. (**H**) The Arnold tongue. The analytically derived PLV is plotted as a function of interaction strength (y-axis) and detuning (x-axis). (**I**) The same as in (H), but for the mean (preferred) phase-relation. Black lines mark the predicted Arnold tongue borders in the noise-free case (ε=|Δ*ω*|).*

To show the applicability of the theory, we first reproduced the three examples shown in Fig.1 and 2 by numerical simulations of equation 1 and by varying detuning Δω and interaction strength ε. We assumed a sinusoidal G(θ) (see Kuramoto model, Breakspear et al., 2010; Kuramoto, 1991) and a phase variability of SD=18Hz. As shown in Fig.3E-G, the same relation between the instantaneous frequency difference modulations and the properties of the phase difference probability distribution were observed as for V1 gamma data. Detuning defined the mean of the frequency modulations, whereas the interaction strength defined the amplitude of the modulations. To obtain a general description of the effect of detuning Δω and interaction strength ε, we mapped the PLV and the mean phase difference (derived analytically) in the Δω-ε parameter space. We observed a triangular synchronization region (Fig.3H) described as the Arnold tongue (Pikovsky et al., 2002). This reflects the fact that stronger interaction strengths ‘tolerate’ larger detuning (Δ*ω<=ε*). Further, a clear phase gradient along the detuning dimension can be observed (Fig.3I). The oscillator with a higher frequency led the oscillator with a lower frequency in terms of their phases.

### Estimating the underlying parameters and function of TWCO in observed data

To demonstrate the underlying principles of V1 gamma synchronization, we aimed to reconstruct its Arnold tongue, a central prediction of the theory. For comparison, we did the same for the coupled PING networks. Further, by estimating the parameters and function of equation 1, we aimed to directly test its accuracy by comparing analytical predictions to experimental observations in V1, and to simulation data from coupled PING networks.

The theory predicts that the phase difference dependent modulations of instantaneous frequency difference (ΔIF(θ)) are determined by the detuning Δω and the interaction term εG(θ). In experimental data, we observed these systematic modulations. Thus, these modulations give information about the detuning and the properties of the interaction term. Specifically, the time-averaged modulation of the instantaneous frequency 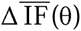 directly relates to the deterministic term Δ*ω*+*ε*G(*θ*), as noise is averaged out (see Supplementary Information). We estimated a single G(θ) function (mutual PRC) and σ value for a given dataset (i.e. each monkey and the PING networks) assuming stability of underlying PRCs and of the noise sources, whereas Δω and ε were estimated for each contact pair and condition. G(*θ*) was estimated by the 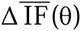 modulation shapes put to unity. The interaction strength ε was estimated by the modulation amplitude of the 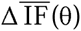. The detuning Δ*ω* was estimated by the average value of the 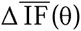 computed over [-π π]. The remaining parameter σ was approximated by finding the σ value for equation 1 that reproduced the observed overall instantaneous frequency variability (see supplementary materials). Given G(θ) and the value σ, the equation can be mathematically (analytically) solved for any values of detuning Δω and interaction strength ε.

### TWCO predicts synchronization properties of weakly coupled PING networks

We first tested the applicability of TWCO for the PING network simulation data. To test for the presence of the Arnold tongue in simulation data, we modulated detuning and interaction strength by varying input drive difference and cross-network connection strength respectively (Fig.4A). From the instantaneous frequency difference modulations (Fig.4B) we reconstructed G(θ), which was approximately a sinusoidal function. This is noteworthy given that the excitatory cross-network connections mainly advanced the phase (Cannon and Kopell, 2015). As discussed later, this was because networks were mutually connected. Further, we estimated the remaining parameters: detuning, interaction strength and the phase noise variance (σ=15Hz). Fig 4C shows for an example level of interaction strength that the analytical predictions of PLV accurately predicted the simulated PLV (model accuracy: R^2^=0.93). Fig 4D demonstrates that mapping the gamma PLV in the Δ*ω* vs. ε parameter space yielded the Arnold tongue with a shape similar to the prediction by the TWCO. Likewise, Fig.4E shows the excellent match between analytical prediction and simulation data for the mean phase difference (model accuracy: R^2^= 0.94), and Fig.4F shows that the mean phase difference of simulated data in the Δ*ω* - ε parameter space yielded the Arnold tongue (Fig.4E-F) with a shape similar to that predicted by the TWCO.

**Fig.4.**
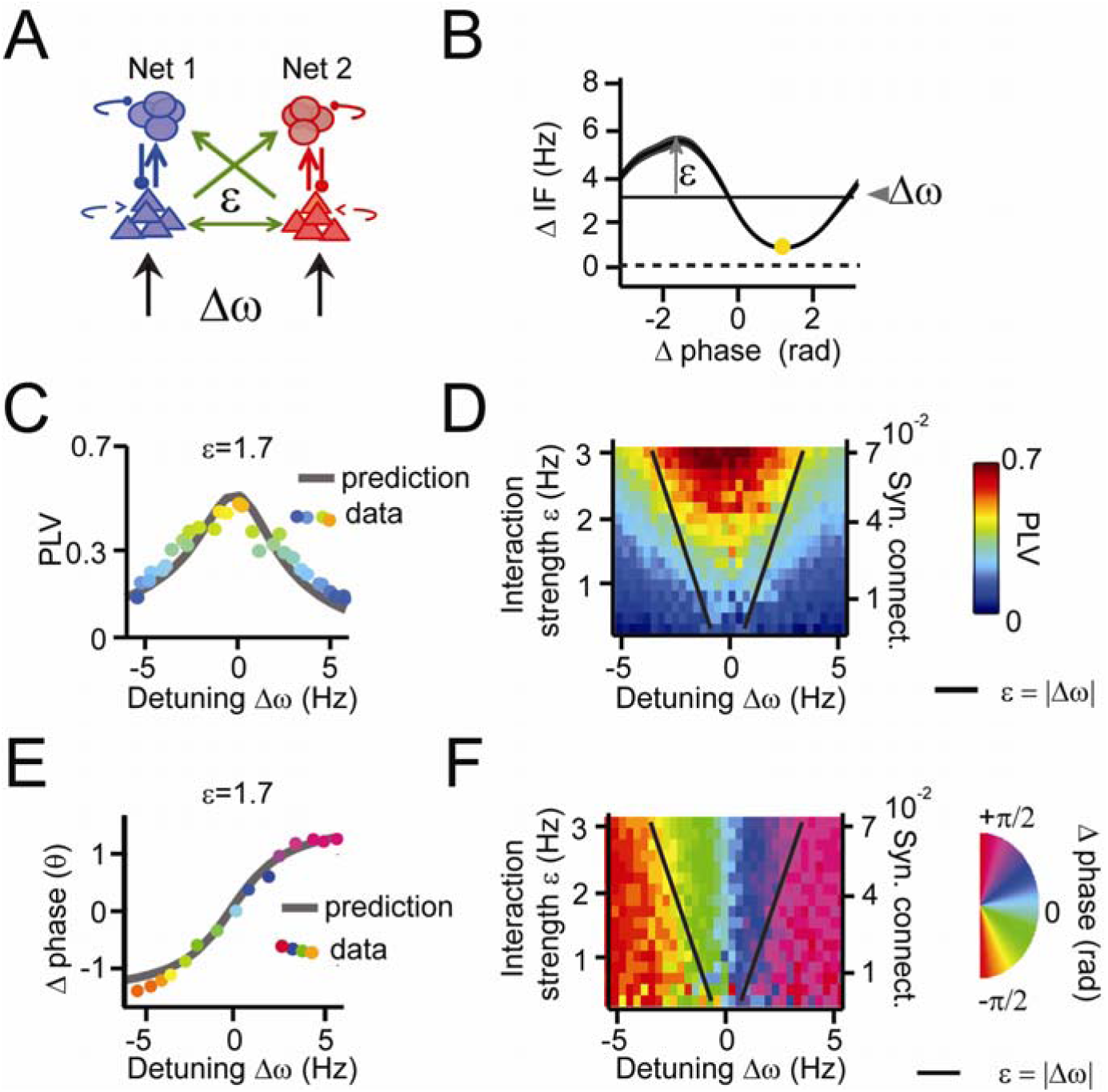
*Applying the theory of weakly coupled oscillators to coupled PING networks. **(A)** Two coupled pyramidal-interneuron gamma (PING) networks (Net 1 and Net 2). Detuning Δω was varied by excitatory input drive, whereas interaction strength ε was varied by inter-network connectivity strength. **(B)** An example plot of averaged phase-dependent modulation of the instantaneous frequency difference (ΔIF) used for estimating ε and Δω. The shape of the modulation indicates the G(θ). **(C)** The simulation PLV at different detuning values Δω (dots colored by PLV) at a single interaction strength value (ε =1.7) was well predicted by the model (gray line). **(D)** The PLV at many interaction strengths and detuning values mapped the Arnold tongue. Black lines mark the predicted Arnold tongue borders in the noise-free case (ε=|Δω|). (**E-F)** As (F-G), but for preferred phase difference θ.*

### TWCO predicts synchronization properties of V1 cortical gamma rhythms

We then tested whether the theory predicted the in vivo data with equal success. In the same manner as with the PING modeling data, we estimated the underlying parameters using the observed modulations of the instantaneous frequency difference (see 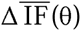 examples in Fig.5A,F), and the phase variance (M1:σ=19Hz, M2:σ=20Hz). The interaction strengths and detuning values were estimated for each channel pair and condition separately. G(θ) was again approximately a sinusoidal function with symmetric negative and positive components (Akam et al., 2012). The interaction strength ε was found to be inversely correlated with the cortical distance between probes (linear regression, M1: R^2^=0.41, M2: R^2^=0.29, both p <10^−10^), in line with V1 horizontal connectivity (Stettler et al., 2002). The detuning Δω was correlated with the contrast difference between probes (linear regression, M1: R^2^=0.31, M2: R^2^= 0.25, both p<10^−10^, Fig.S2). Combining gamma PLV estimates from all recorded V1 pairs, we were able reconstruct the Arnold tongue as a function of Δω and ε in both M1 and M2 (Fig.5C/G) confirming a central theoretical prediction. To better evaluate the accuracy of the theory, we derived analytical predictions for different Δω and ε by solving equation 1 using the estimated parameters. We found that the gamma PLV variation over all single contact pairs were substantially captured by the analytical predictions as a function of Δω and ε (model accuracy: M1: R^2^=0.18, n=7245, M2: R^2^= 0.32, n=7938). The observed population means for different Δω and ε values followed the analytical predictions well (model accuracy: M1: R^2^=0.83, M2: R^2^= 0.86, both n=638). In Fig.5D/H we plotted a horizontal cross-section of the Arnold tongue that shows the good fit between the prediction and observed population means. The observation of a gamma Arnold tongue across the V1 middle-superficial layers was confirmed also for deep layer contacts (Fig.S4). We then mapped the mean phase difference (preferred phase-relation) between V1 gamma rhythms as function of Δω and ε. We observed a clear phase gradient in both monkeys across the detuning dimension (Fig.5E/I). The phase spread (see also Fig.5F/J) had a range of nearly –pi/2 to pi/2 in both M1 and M2, as predicted by the shape of G(θ). Gamma rhythms with the higher frequency of a pair had the leading preferred phase relation. The mean phase difference increased with increased detuning. For given detuning, stronger interaction strength led to a reduction of the phase difference. Over all single contact pairs the mean phase difference was substantially captured by the analytical predictions (model accuracy: M1: R^2^=0.56, n=7245, M2: R^2^=0.3, n=7938). The observed population means for different Δω and ε values followed the analytical predictions precisely (model accuracy: M1: R^2^=0.92, M2: R^2^=0.88, both n=638).

**Fig.5.**
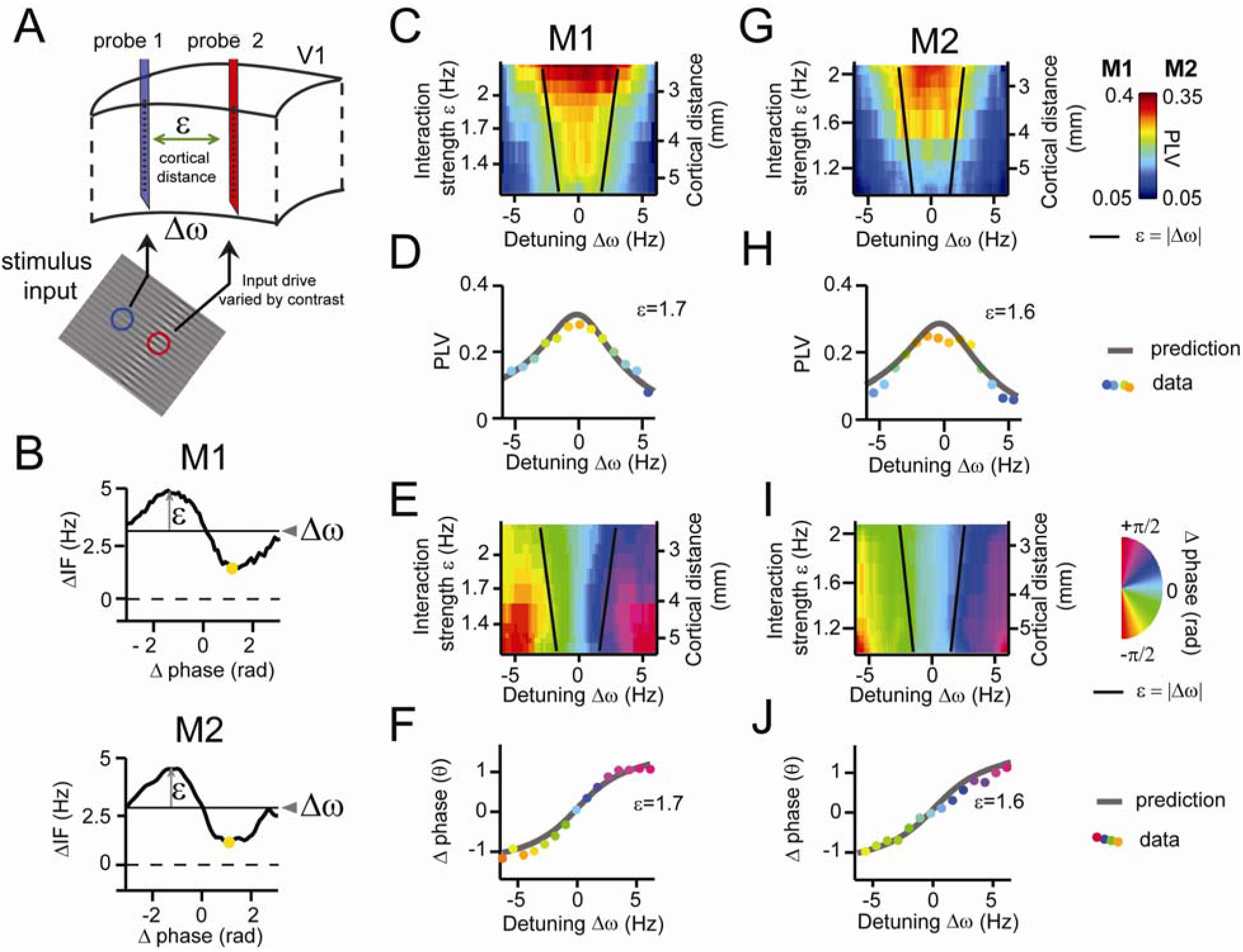
*Predicting V1 gamma synchronization. **(A)** Illustrative schema showing how detuning Δω and interaction strength ε of V1 gamma relate to local stimulus contrast and cortical distance respectively. (**B)** An example plot of averaged phase-dependent modulation of the instantaneous frequency difference (ΔIF) used for estimating ε and Δω for monkey M1 (top) and M2 (bottom). The shape of the modulation indicates the G(θ). **(C-F)** Results from M1. (**C)** Observed PLV (dots) and analytical prediction (gray line) as a function of detuning Δω for one level of interaction strength (ε=1.7). (**D)** Combining different detuning Δω and interaction strengths ε we observed a triangular region of high synchronization, the Arnold tongue. Black lines mark the predicted Arnold tongue border as expected from the noise-free case (ε=|Δω|) (**E)** Analytical prediction (gray) and experimentally observed preferred phase differences (dots colored by phase difference) as a function of detuning Δω for one level of interaction strength (ε=1.7). (**F)** Similar to D), but now plotting the preferred phase difference. **(G-J)** As (C-F) but for M2 population data. Color coding of dots in C, H, E, I is as indicated in color scales in panels just below them.*

We confirmed the PLV and phase difference analysis in spike-CSD (spike-field) and spike-spike measurements (Fig.S5).

### Comparison of TWCO, PING and V1 gamma synchronization

To reveal the individual contributions of detuning and interaction strength in regulating the PLV and the mean phase difference, we applied a multiple regression approach with detuning, interaction strength and amplitude as factors (Fig.6). The contributions were expressed in explained variance (R^2^). We found that the TWCO (Fig.6A) reflected the same pattern of contributions as we observed for PING (Fig.6B) and V1 gamma rhythms (Fig.6C). The phase locking value (PLV) was mainly determined by interaction strength and more weakly by detuning. The mean phase difference was however primarily determined by detuning and only weakly by interaction strength. Interaction strength affected the mean phase difference through an interaction effect with detuning by changing the detuning-to-phase-difference slope (interaction effect in Fig.6). In addition to the predictions of TWCO, we observed weak effects of the oscillation amplitude on the PLV and on the mean phase difference in PING and V1 gamma data. Amplitude differences between gamma rhythms can lead to asymmetric interaction strengths that shift the precise PLV and the preferred phase-relation. Further, in both PING and V1 data, we observed phase-dependent instantaneous amplitude modulations (Fig.S6). However, the analytical predictions and multiple regression analysis are in agreement in showing that detuning and interaction strength (frequency modulations) represent the main parameters for regulating V1 gamma synchronization.

**Fig.6:**
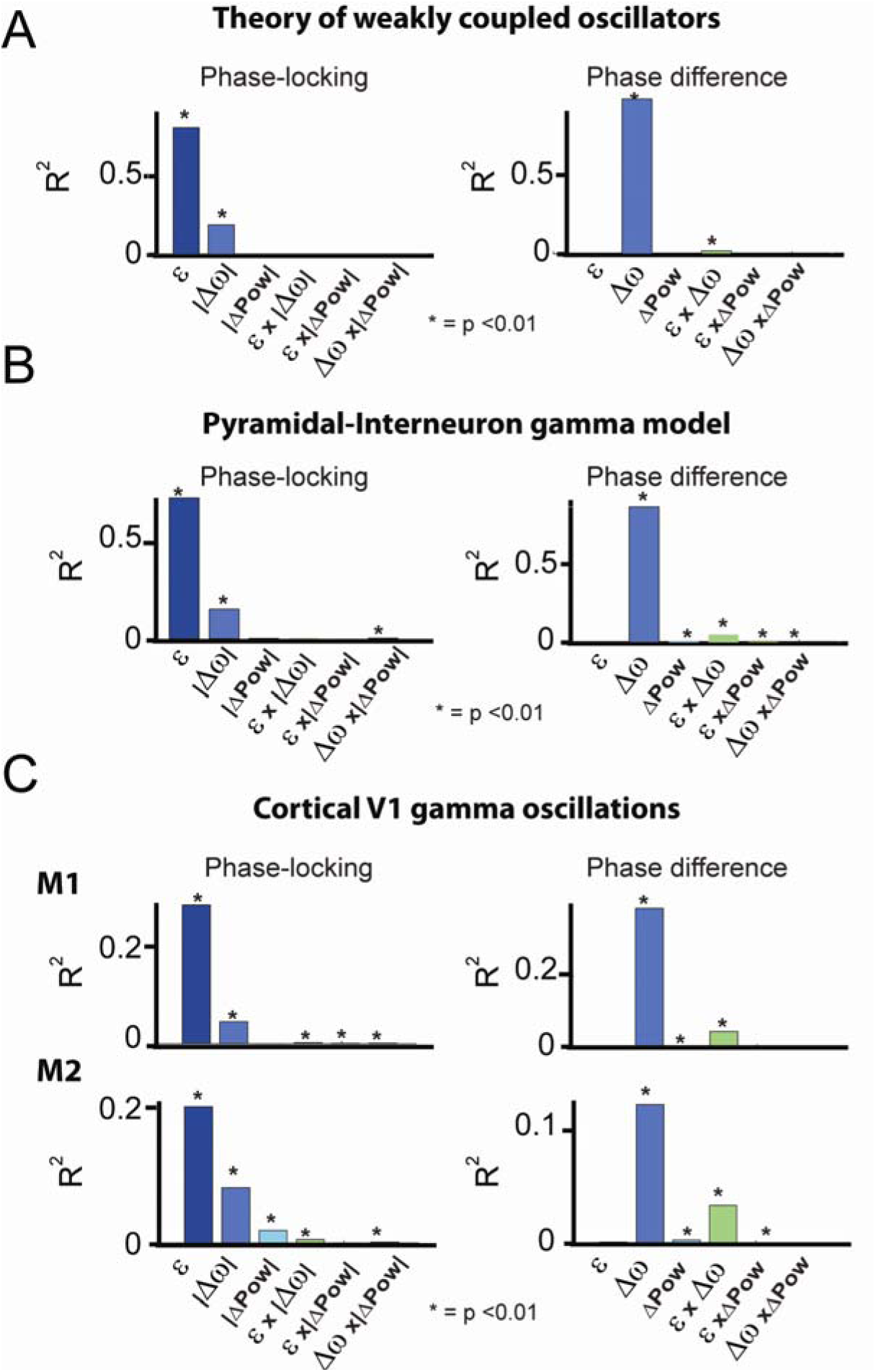
*Multiple regression analysis of PLV and mean phase difference. (**A**) TWCO numerical simulations (n=673) including different detuning (-6Hz to 6Hz) and interaction strengths (0< ε < 3.5). (**B**) PING network simulations (n=697) including different inter-network connection strengths (0.008–0.072) and input drive differences (−5 to 5). (**C**) Macaque V1 single contact data including all contact pairs and conditions (M1: n=7245, M2: n=7938). A significance value below P<0.01 is marked with an asterisk. The results for PLV are on the left and for phase difference are on the right. Contributions are expressed in explained variance (R^2^).*

## DISCUSSION

The present study shows that gamma synchronization in awake monkey V1 adheres to theoretical principles of weakly coupled oscillators (Ermentrout and Kleinfeld, 2001; Hoppensteadt and Izhikevich, 1998; Kopell and Ermentrout, 2002; Kuramoto, 1991; Pikovsky et al., 2002; Winfree, 1967), thereby providing insight into the synchronization regime of gamma rhythms and its principles. Given the generality of the synchronization principles, they are likely to apply to other brain regions and frequency bands.

### Intermittent synchronization: The role of non-stationary frequency modulations

Our findings reveal the importance of phase-dependent frequency modulations for synchronizing V1 gamma rhythms. The same modulations were observed in a general biophysical model of gamma rhythms. These modulations show that a fixed and common frequency is not required for phase coordination. To the contrary, stronger non-stationary frequency modulations led to stronger synchronization, and thus to more reliable phase coordination. These modulations arise naturally in the intermittent synchronization regime (Izhikevich, 2007; Pikovsky et al., 2002), when oscillators cannot remain in a stable equilibrium due to detuning and noise. Given the variable nature of gamma rhythms in vivo (Atallah and Scanziani, 2009; Burns et al., 2010; Ray and Maunsell, 2010; Roberts et al., 2013), intermittent synchronization is the most likely regime for their phase coordination. Although complete synchronization is not achieved in this regime, phase coordination remains sufficiently robust to influence the strength and directionality of information flow (Battaglia et al., 2012; Buehlmann and Deco, 2010; Fries, 2015; Maris et al., 2016), by rendering particular phase-relations more likely than others. The observation of non-stationary frequency modulations also has methodological implications. Gamma rhythms are often studied with stationary methods, for example spectral coherence or stationary granger measures, yet our findings are not in line with the (weak-sense) stationarity assumption (Lachaux et al., 1999; Lowet et al., 2016). Time-resolved non-stationary methods are therefore more appropriate to study the dynamics underling gamma synchronization.

### The interaction function of V1 gamma rhythms

We show that the shape of the frequency modulations reflects the underlying interaction function, the PRC (Hoppensteadt and Izhikevich, 1998; Kopell and Ermentrout, 2002; Kuramoto, 1991; Pikovsky et al., 2002; Winfree, 1967). The PRC defines how the oscillators advance or delay each other’s phase development to coordinate their phase-relation. We observed symmetric sinusoidal-like functions in both PING and in V1 gamma that resemble the basic function of the widely-used Kuramoto-model (Breakspear et al., 2010). This is in agreement with the biphasic PRC of gamma rhythms observed in the rat hippocampus (Akam et al., 2012). In agreement with our symmetric G(θ), we observed symmetric Arnold tongues (Izhikevich, 2007; Kopell and Ermentrout, 2002; Pikovsky et al., 2002). Importantly, here we estimated the mutual (bidirectional) PRC, the G(θ). This function can be symmetric (equal magnitude of phase advance and delay), despite asymmetric individual (unidirectional) PRCs, as long as the rhythms interact approximatively equally strongly, which is a plausible assumption between V1 locations. Therefore, our results are not per se at odds with other studies that have indicated asymmetric individual PRC in neural data (Cannon and Kopell, 2015; Wang et al., 2013). Unidirectionally connected neural groups, for example between certain cortical areas, might have asymmetric PRC and hence an asymmetric Arnold tongue. In this situation a frequency difference between cortical areas (Bosman et al., 2012; Cannon et al., 2014) might be favorable for optimal information transmission. This hypothesis could be tested between gamma rhythms recorded from unidirectionally connected cortical areas.

### The Arnold tongue and the regulative parameters of gamma synchronization

Previous studies have established diversity in the phase-locking (Eckhorn et al., 2001; Gray and Singer, 1989; Ray and Maunsell, 2010) and in the phase-relation (Maris et al., 2016; Vinck et al., 2010) of gamma rhythms in the primate visual cortex. However, how this diversity is regulated was not well established. Here, we show that two parameters mainly determined gamma synchronization: the detuning (frequency difference) and the interaction strength ε (frequency modulations). This was highlighted in the mapping of the Arnold tongue, offering a graphical understanding of how these parameters shape gamma-band synchronization. Detuning represents a desynchronization force, whereas the interaction strength represents a synchronization force. The former was modulated by input drive differences, and the latter by connectivity strength. Their interplay defined the resultant phase-locking strength and the preferred phase-relation between gamma rhythms. The observed role of detuning is in agreement with a previous study in the rat hippocampus (Akam et al., 2012), in which optogenetic entrainment strength and phase of gamma rhythms were dependent on the frequency-detuning. The results also agree with theoretical conceptions on oscillatory interactions (Ermentrout and Kopell, 1984; Hoppensteadt and Izhikevich, 1998; Sancristóbal et al., 2014; Tiesinga and Sejnowski, 2010). We suggest that small detuning values (mainly <Δ10Hz) reported in the present study and much larger shifts in the gamma frequency-range (25-50Hz to 65-120Hz) reported in the rat hippocampus and cortex (Colgin et al., 2009) represent different but complementary mechanisms for controlling gamma synchronization. In this perspective, large shifts in the frequency-range could selectively turn on or off gamma-mediated information flow between brain regions, whereas fine frequency detuning modulates the exact strength and direction of the gamma-mediated information flow. The role of instantaneous frequency modulations, defining the interaction strength, reflects the overall ability of two cortical locations to engage in gamma-band synchronization. These modulations are mediated by anatomical connectivity and further modified by oscillation amplitude. Hence, an important source of instantaneous V1 gamma frequency modulations (Bosman et al., 2009; Burns et al., 2011, 2010; Roberts et al., 2013) is the underlying network (intermittent) synchronization process. Instantaneous gamma frequency fluctuations have also been observed in the rat hippocampus by Atallah and Scanziani (2009). Their data suggested that these fluctuations, which reflected rapid phase shifts due to changes in excitation-inhibition balance, might be critical for gamma-mediated information flow. In line with this notion, we show that these cycle-by-cycle modulations are essential for regulating synchronization properties between gamma rhythms.

### Role of V1 gamma synchronization for visual processing

In our experiment, detuning was dependent on the local contrast difference (Ray and Maunsell, 2010; Roberts et al., 2013), known to change neural excitation in V1 (Sclar et al., 1990), while the interaction strength was dependent on the underlying horizontal connectivity strength, here varied by cortical distance (Stettler et al., 2002). Gamma synchronization is therefore informative about the sensory input (Besserve et al., 2015) and informative about the underlying structure of connectivity. Indeed, the frequency of gamma rhythms is modulated by various sensory stimuli (Fries, 2015) and by cognitive manipulations (Bosman et al., 2012; Buzsáki and Wang, 2012; Fries, 2015) suggesting that frequency control is critical for functional V1 gamma-band coordination. The horizontal connectivity in V1 is not only local, but also exhibits remarkable tuning to visual features, orientation being a prime example (Stettler et al., 2002). Hence, innate and learned connectivity patterns likely affect the interaction strength and hence the synchronization patterns of gamma rhythms within V1. These properties suggest V1 gamma as a functional mechanism for early vision (Eckhorn et al., 2001; Gray and Singer, 1989) by temporally coordinating local neural activity as a function of sensory input and connectivity. However, in line with previous studies (Eckhorn et al., 2001; Palanca and DeAngelis, 2005), V1 gamma synchronization was found to be mainly local and hence not likely to ‘bind’ whole perceptual objects. Furthermore, recent studies on the gamma-band response during natural viewing (Brunet et al., 2013; Hermes et al., 2014) have found variable levels of synchronization power for different natural images. In line with these observations, the revealed Arnold tongue of V1 gamma implies that natural image parts with high input/detuning variability (heterogeneity) will induce no or weak synchronization, whereas parts with low input/detuning variability (homogeneity) will induce strong synchronization. This is also in line with proposals linking gamma synchronization with surround suppression/normalization (Gieselmann and Thiele, 2008; Ray et al., 2013) and predictive coding (Vinck and Bosman, 2016). Our findings and interpretation shed new light onto the operation of gamma synchronization in the brain and will permit new and more detailed description of the mechanisms by which synchronization is regulated by cognitive and sensory inputs.

### Experimental Procedures

#### Species used and surgical procedures

Two adult male rhesus monkeys were used in this study. A chamber was implanted above early visual cortex, positioned over V1/V2. A head post was implanted to head-fix the monkeys during the experiment. All the procedures were in accordance with the European council directive 2010/63/EU, the Dutch ‘experiments on animal acts’ (1997) and approved by the Radboud University ethical committee on experiments with animals (Dier-Experimenten-Commissie, DEC).

#### Recording methods

V1 recordings were made with 2 or 3 Plexon U-probes (Plexon Inc.) consisting of 16 contacts (150µm inter-contact spacing). We recorded the local field potential (LFP) and multi-unit spiking activity (MUA). For the main analysis we used the current-source density (CSD, (Vaknin et al., 1988)) to reduce volume conduction. We aligned the neural data from the different laminar probes according to their cortical depth and excluded contacts coming from deep V2. Layer assignment was based on the stimulus-onset CSD profile (Schroeder et al., 1991) and the interlaminar coherence pattern (Maier et al., 2010). Receptive field (RF) mapping was achieved by presenting at fast rate high-contrast black and white squares pseudorandomly on a 10x10 grid (Roberts et al., 2013). For RF mapping we used CSD signals and spikes.

#### Task and visual stimuli

The monkeys were trained for head-fixation and were placed in a Faraday-isolated darkened booth at a distance of 57cm from a computer screen. Stimuli were presented on a Samsung TFT screen (SyncMaster 940bf, 38°x30° 60Hz). During stimulation and pre-stimulus time the monkey maintained a central eye position (measured by infra-red camera, Arrington, 60Hz sampling rate). The monkey´s task was to passively gaze on a fixation point while a stimulus was shown. The monkey was rewarded for correct trials. The local stimulus contrast was manipulated in a whole-field static square-wave grating (2 cycles/degree, presented at two opposite phases randomly interleaved). Contrast was varied smoothly over space such that different RFs had different contrast values. The direction of the contrast difference was parallel to the arrangement of RFs and orthogonal to the orientation of the grating. The stimulus was isoluminant at all points and was isoluminant with the pre-stimulus grey screen. We presented 9 different contrast modulation conditions (Table.S1). Cortex software (http://dally.nimh.nih.gov/index.html) was used for visual stimulation and behavioral control.

#### Data analysis

To investigate dynamical changes in the gamma phase and frequency over time we estimated the instantaneous gamma phase and frequency using the singular spectrum decomposition of the signal (SSD (Bonizzi et al., 2014), see https://project.dke.maastrichtuniversity.nl/ssd/) combined with Hilbert-Transform or wavelet-decomposition. The phase-locking value (PLV) was estimated as the mean resultant vector length (Lachaux et al., 1999) and the preferred phase-relation as the mean resultant vector angle. For experimental data, we estimated the signal-to-noise ratio (SNR) to reduce the influence of measurement noise on estimates. Phase flipping due to CSD computation was corrected.

#### Theoretical and computational modelling

Using the theory of weakly coupled oscillators we investigated the phase-locking as well as the mean phase difference of two mutually coupled noisy phase-oscillators with variable frequency difference (detuning) and interaction strength. The stochastic differential equation was solved analytically (Pikovsky et al., 2002). The analytical results correctly predicted the numerical simulations. In addition, we simulated two coupled excitatory-inhibitory spiking networks generating gamma oscillations using the Izhikevich-type neuronal model (Izhikevich, 2003). The detuning between the networks was altered by changing the difference in excitatory input drive. The interaction strength was altered by changing the cross-network synaptic connection strength.

#### Statistics

The accuracy of the theoretical predictions for the experimental data was quantified as the explained variance R^2^. In addition, to evaluate the contribution of different parameters we used a multiple regression approach (Matlab function fitlm, The MathWorks Inc.).

## Acknowledgements

### Author contributions

E.L., M.J.R. and P.DW. designed the experiment. E.L. and M.J.R. conducted the recordings. Data analysis and the writing of the manuscript was by E.L. with support by M.J.R, A.P., B.G. and P.DW.

### Acknowledgments

We thank N.Kopell, W.Singer, A.Bastos, P.Fries, C.Micheli, F.Smulders, J.v.d.Eerden, J.Karel, P.Bonizzi, A.Hadjipapas, A.A.v.d.Berg. Supported by NWO VICI grant 453-04-002 to PDW and NWO VENI grant 451-09-025 to MJR. All data are stored at the Department of Psychology and Neuroscience, Maastricht University, The Netherlands. We thank the Radboud University Nijmegen for hosting our experiments, and staff of the Central Animal Facility (CDL) for expert assistance.

